# Global Footprint of the Multidrug Resistance Island Ec17R and Resistance Gene Co-Occurrence in Pathogenic *Escherichia coli* Isolates

**DOI:** 10.1101/2025.10.10.681733

**Authors:** Elijah R. Bring Horvath, O’Connor J. Matthews, William J. Brazelton, Edgar J. Hernandez, Hillary Crandall, Sara M. Lenherr, Jaclyn M. Winter, Matthew A. Mulvey

**Affiliations:** Department of Pharmacology and Toxicology, University of Utah, Salt Lake City, Utah, 84112; School of Biological Sciences, University of Utah, Salt Lake City, Utah, 84112; Henry Eyring Center for Cell and Genome Science, University of Utah, Salt Lake City, Utah, 84112; Department of Biomedical Informatics, University of Utah, Salt Lake City, Utah, 84112; Department of Pediatrics, University of Utah, Salt Lake City, Utah, 84112; Department of Surgery, Genitourinary Injury and Reconstructive Urology, University of Utah, Salt Lake City, Utah, 84112

**Keywords:** antibiotic resistance, Bayesian nets, uropathogenic, UTI, ExPEC

## Abstract

Multidrug resistant (MDR) bacterial pathogens are a major threat to global health, limiting treatment options for common infections. Extraintestinal Pathogenic *Escherichia coli* (ExPEC), a leading cause of bloodstream and urinary tract infections (UTIs), are often resistant to one or more antibiotic classes. Previously, we identified a clonal group of ExPEC strains (P1A) that persisted over a 5-year period from 2012 to 2016 within a female patient who suffered from frequent recurrent UTIs. A subset of these isolates carried a plasmid-borne MDR island (Ec17R) harboring 17 resistance genes. Sampling of fecal and urine samples in 2019 indicated that the patient remained colonized with the P1A lineage, including Ec17R-positive strains, over 7 years after collection of the first P1A isolates in 2012. Highlighting the public health relevance and mobility of Ec17R, we found that Ec17R-like islands are globally distributed across diverse bacterial species from various environmental, agricultural, and clinical sources, including multiple ExPEC isolates from local pediatric patients. By applying clustering approaches and Bayesian network modeling to 267 pediatric ExPEC isolates, we found that functionally distinct classes of resistance genes (including several heavy metal resistance genes) have a high probability of co-occurrence, possibly reflecting carriage within MDR islands like Ec17R. Finally, we observed that strains within the P1A lineage are recalcitrant to antibiotics for which they have no known resistance mechanisms, suggesting that these pathogens have means beyond their formidable array of resistance genes to survive within a host for years despite the administration of numerous, robust antimicrobial treatments.

**SIGNIFICANCE:** Multidrug resistance (MDR) in bacteria is a growing global health crisis that compromises our ability to treat routine infections. In this study, we investigated a clonal lineage of pathogenic *Escherichia coli* that persisted for many years in a patient with recurrent urinary tract infections. A subset of the *E. coli* strains carried by this patient possessed a large genomic island encoding resistance to multiple antibiotic classes. This MDR island is globally distributed across diverse bacterial species and niches, from clinical samples to agricultural and environmental reservoirs. Using probabilistic modeling of nearly 300 clinical isolates, we identified networks of resistance gene co-occurrence that link antibiotic and heavy metal resistance, suggesting the potential for environmental pollutants to contribute to MDR dissemination. Notably, patient-derived isolates also survived certain clinically relevant antibiotic treatments despite lacking known resistance mechanisms, highlighting tolerance and persistence as important, often overlooked drivers of therapeutic failure.

## INTRODUCTION

Antibiotic resistance (AR) is currently recognized by the World Health Organization as an urgent global health threat (1, 2). Its continued rise reflects decades of suboptimal antibiotic stewardship, including antibiotic overuse, inappropriate use of antibiotics for non-bacterial infections, and extensive agricultural use, the latter of which serves as a route of exposure of antibiotics and the associated selective pressures to environmental bacteria (3–5). Additionally, there has been a marked decline in the development of novel antibiotics since the “Golden Age” of antibiotic discovery during the 1940s– 1960s (3, 6–9). This problem is compounded by the fact that most anti-infectives are naturally occurring (i.e., natural products) or based on naturally occurring chemical scaffolds, so intrinsic resistance mechanisms often exist in nature long before a drug is introduced clinically (3, 7, 10–12). Recently, the CDC estimated that nearly 3 million AR infections and >35,000 associated deaths occur annually in the United States, representing a >150% rise in expected mortality in just over six years (1, 13–15). Although infection control measures and stewardship have driven important improvements in some settings (15), the overall burden of AR microbes continues to expand.

Within this crisis, multidrug-resistant (MDR) bacteria pose the greatest clinical challenge (5, 13). MDR bacterial pathogens are frequently resistant to multiple antibiotic classes, leaving few effective treatment options. Previously, we analyzed a clonal population (P1A) of sequence type 131 (ST131) Extraintestinal Pathogenic *Escherichia coli* (ExPEC) isolates from a patient (P1) at the University of Utah hospital who experienced chronic, recurrent urinary tract infections (rUTI) dating back to the early 1970s (16). Phylogenomic and plasmid analyses of urine and fecal isolates collected between 2012– 2016 revealed a subset of the PA1 strains harboring a plasmid-borne MDR island (designated here as Ec17R) containing 17 genes that are associated with resistance to multiple clinically important classes of antibiotics, including aminoglycosides, macrolides, tetracycline, sulfonamides, and trimethoprim (16). Notably, these MDR strains persisted year-to-year despite the administration of frequent and diverse antibiotic treatments.

MDR determinants are frequently shared among bacteria through horizontal gene transfer, most often via plasmids, transposons, and integrative genomic islands. These elements enable the linkage of multiple resistance genes into larger assemblies that can move between strains and even across species boundaries (17). Importantly, MDR islands can contain not only antibiotic resistance genes (ARGs), but also heavy metal resistance genes (HMRGs). If these determinants are physically clustered, selective pressures unrelated to antibiotic use—such as environmental exposure to heavy metal pollutants—can drive the co-selection and maintenance of ARGs (18–20). In addition, genomic proximity means that transfer events initiated under environmental stress may disseminate both HMRGs and ARGs together, broadening the ecological and clinical contexts in which multidrug resistance can emerge and persist.

Canonical ARGs explain survival under many treatments, yet bacteria may also withstand antibiotic exposure in the absence of known resistance mechanisms (21–23). The cellular processes underlying such antibiotic recalcitrance—often manifesting as tolerance or persistence-like phenotypes—remain poorly understood, but these processes can enable regrowth to infectious population sizes and drive cycles of treatment and recurrence, or recrudescence (21, 22). Together, resistance and recalcitrance constitute a dual barrier to effective therapy.

As AR rates are projected to increase among ExPEC strains and other pathogens (3, 6, 15), robust stewardship must be coupled to deeper understanding of MDR architectures and their epidemiology. Here, we characterize the extensive MDR genomic island Ec17R, evaluating its global distribution and origins as well as those of key resistance modules within Ec17R. We then analyze co-occurrence patterns among ARGs and HMRGs across 267 ExPEC isolates from hospitalized pediatric patients at Primary Children’s Hospital (Salt Lake City, Utah) using a discrete Bayesian network structure learning probability model. Finally, we provide evidence that the P1A rUTI isolates are able to survive exposure to ampicillin and ertapenem at unexpectedly high levels despite having *in vitro* sensitivity and a lack of any known resistance mechanisms to these antibiotics, underscoring the clinical relevance of antibiotic recalcitrance mechanisms alongside AR.

## RESULTS

### Multidrug Resistant Genomic Island in rUTI-Causing *Escherichia coli*

In our 2019 publication (16), we reported the persistence of the P1A ExPEC lineage within rUTI patient P1 from 2012 to 2016. Over the next three years, patient P1 continued to experience frequent rUTIs, despite receiving culture-specific antibiotic treatments which did not provide any lasting relief. In 2019, patient P1 provided additional fecal and clean-catch urine samples that were plated on selective media (MacConkey agar) with or without either ciprofloxacin or trimethoprim. At the time of collection, the patient was asymptomatic for UTI and had not received any antibiotics for the preceding three weeks. Bacteria present in the patient’s urine were sparse, and so centrifugation was used to concentrate the sample prior to plating. In the end, only 12 colonies were recovered from the urine sample. These bacteria, as well as 24 fecal isolates from patient P1, were analyzed by whole-genome sequencing. Consistent with our previous findings, all strains were classified as ST131 (**Figure 1A**) and exhibited greater genomic similarity to one another and to earlier isolates from this patient than to other *E. coli* strains (**Figure 1**), indicating the continued persistence of the P1A lineage in this patient from at least 2012 into 2019. Among the sequenced isolates, 12 carried MDR regions identical or highly similar to Ec17R (**Figure S1**), which was originally described in ExPEC strain U15A (**Figure 2A**) (16). For clarity, ‘Ec17R’ describes MDR islands with identical ARG compliments as that described in U15A (Example: **Table 1**, plasmids pKF3 and p283149-FII); ‘Ec17R-like’ describes regions that differ by one or more ARGs but encode most (≥50%) of the ARGs found in Ec17R (Example: **Table 1**, plasmids P1-09-02E and pCf75). Notably, trimethoprim resistance appeared to serve as a marker for Ec17R/Ec17R-like regions in this population. Indeed, as in our previous study (16), all PA1 isolates recovered from trimethoprim-selective media harbored the MDR island.

**Figure 1.**
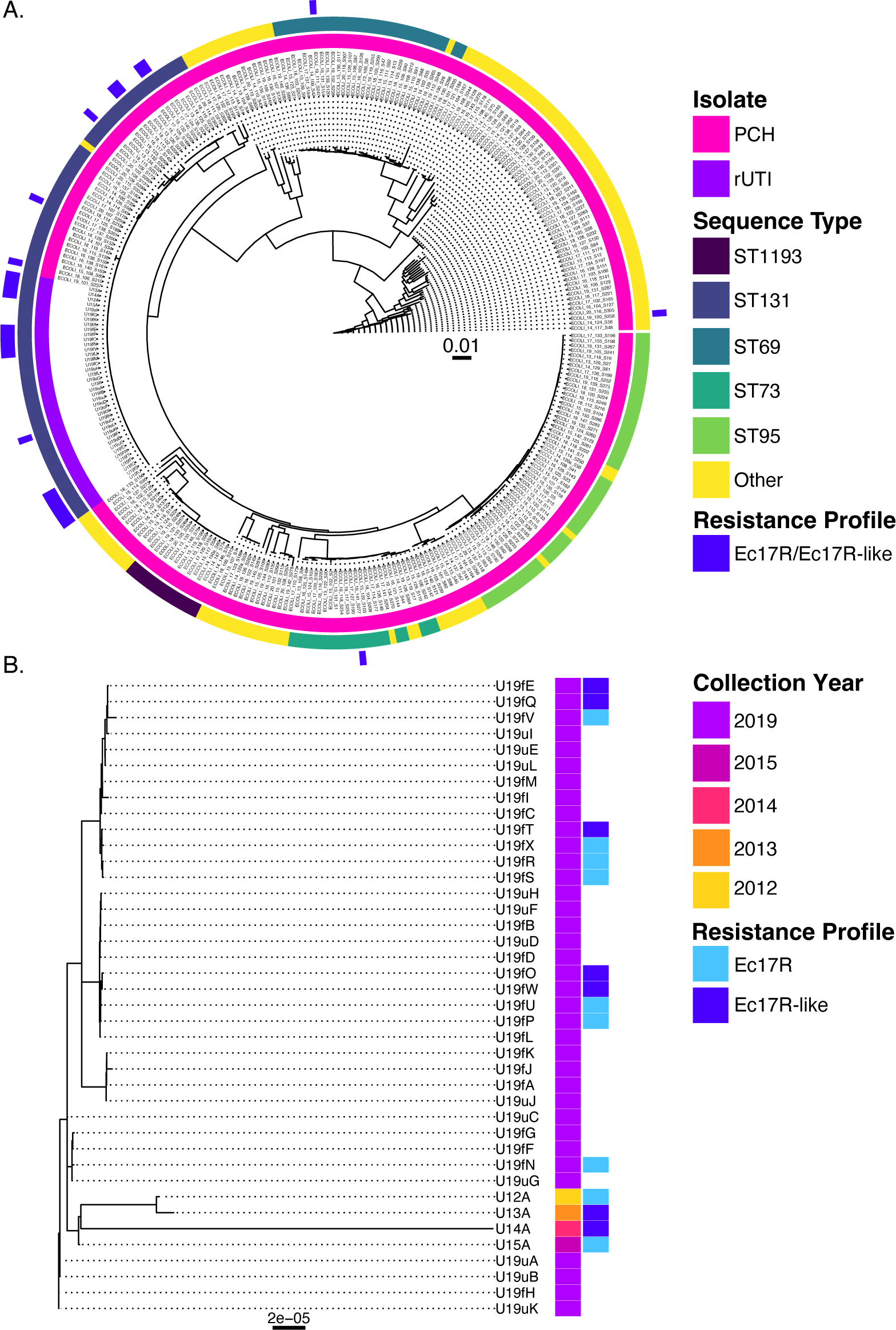
Maximum likelihood (ML) phylogenetic trees highlight persistence of the PA1 ExPEC lineage within patient P1 for at least 8 years. (**A**) ML tree of all and sequenced PA1 (rUTI) and PCH patient isolates. The first (innermost) ring denotes isolate source (Primary Children’s Hospital or rUTI patient P1, PA1); second ring (middle) indicates sequence type; third ring (outermost) indicates strains carrying Ec17R or an Ec17R-like multidrug resistance (MDR) island. (**B**) ML tree of rUTI patient P1 isolates collected between 2012 and 2019, with collection years and presence of Ec17R/Ec17R-like MDR regions indicated.

**Figure 2.**
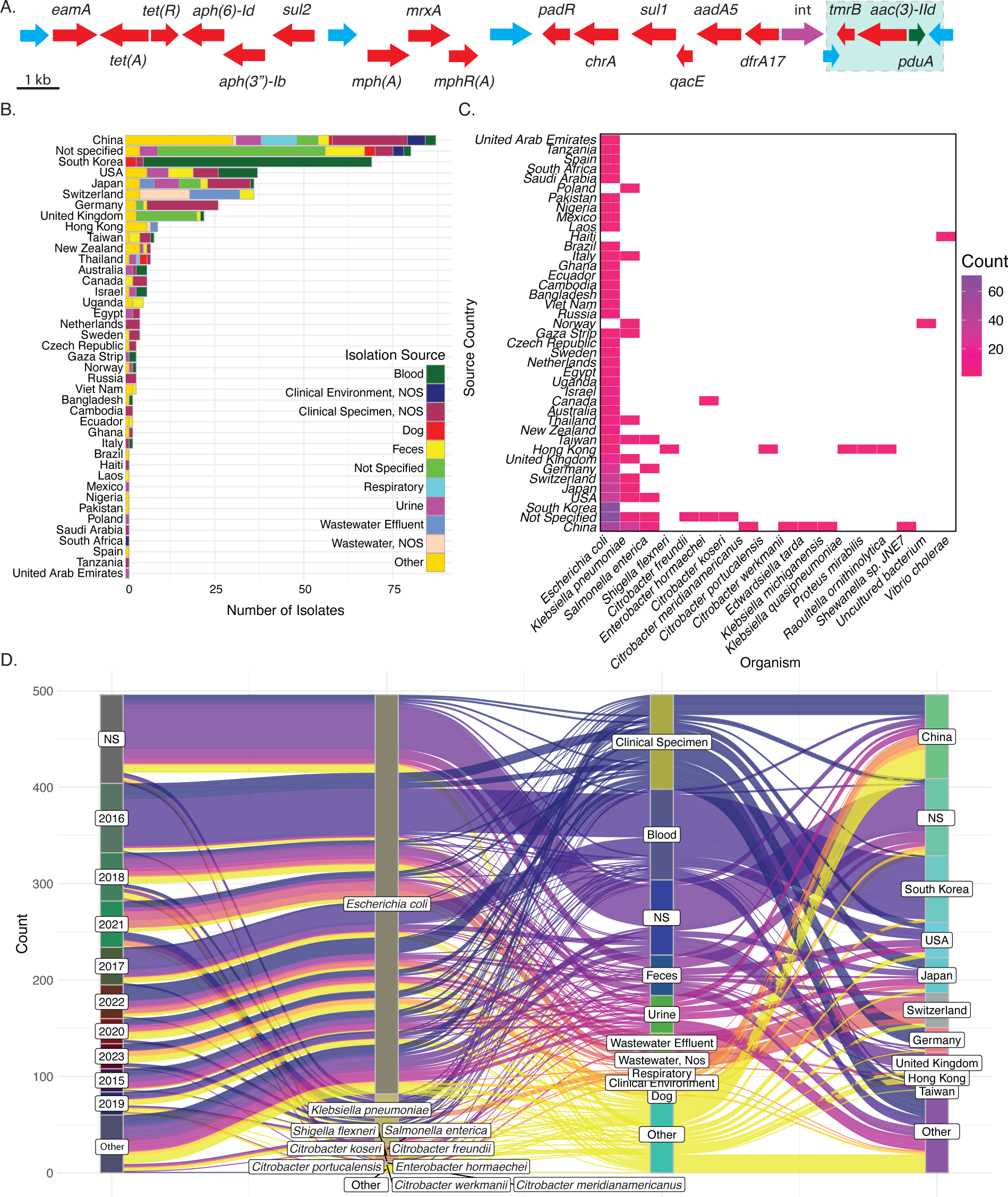
Global footprint of Ec17R and Ec17R-like multidrug resistance islands. (**A**) Ec17R identified in plasmid pU15A_A harbored by *E. coli* strain U15A, recovered from rUTI patient P1. Arrows represent open reading frames and point in the direction of transcription. Red, predicted resistance genes. Blue; transposable elements. Green; predicted non-resistance genes. (**B-D**) Sources of strains harboring Ec17R-like resistance islands identified via NCBI nucleotide BLAST searches. (**B**) Most common sources of strains with Ec17R-like islands and associated countries. (**C**) Distribution of species harboring Ec17R-like islands and associated countries of origin. (**D**) Years of isolation for strains with Ec17R-like islands linked with associated microbes, sources, and countries of origin. The green box in panel A indicates the variable region of Ec17R in the P1A population. NS: not specified.

**Table 1.**
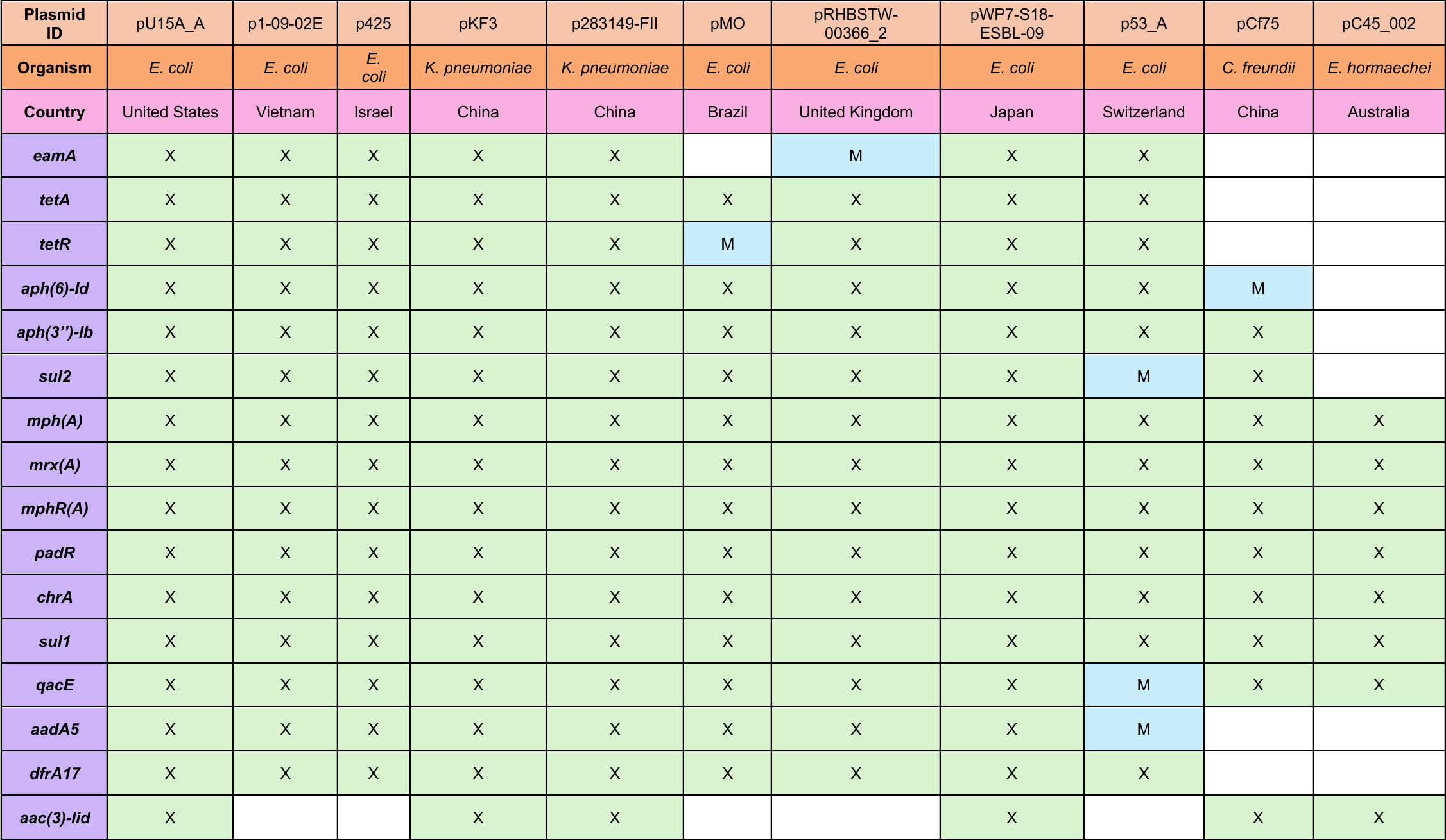
Ec17R-like multidrug resistance islands identified in strains from across the globe. The table indicates plasmids from various isolates (*E. coli, Klebsiella pneumoniae, Citrobacter freundii,* and *Enterobacter hormaechei*) that possess regions of genes that are highly similar or exact matches with those in Ec17R. ‘X’ denotes an exact nucleotide match to genes within Ec17R. ‘M’ signifies that the same gene was found, but did not have an exact nucleotide match. pKF3, pCf75, and pC45_002 are examples of high-coverage, middle-coverage, and low-coverage matches, respectively (minimum ≥85% query coverage, ≥99% sequence identity). *eamA,* small molecule efflux; *tetA/R*, tetracycline resistance efflux operon; *aph*(*6*)*-Id/aph(3’’)-Ib*, streptomycin resistance O-phosphotransferase; *sul2*, sulfonamide resistance; *mph(A)*, macrolide 2’-phosphotransferase; *mrx(A)*, macrolide efflux; *mphR(A)*, macrolide resistance cassette transcriptional regulator; *padR/chrA*, chromate resistance efflux operon; *sul1*, sulfonamide resistance; *qacE*, quaternary ammonium efflux; *aadA5*, aminoglycoside adenylyl-transferase; *dfrA17*, trimethoprim resistance; *aac*(*3*)*-IId*, aminoglycoside resistance acetyltransferase.

Two primary sources of variation were observed within Ec17R-like regions carried by the sequenced PA1 strains that were collected between 2012 and 2019. First, the aminoglycoside resistance gene *aac*(*3*)*-IId* was variably present (**Figure 2A, green box**). This gene occurred within a small gene cluster that also encoded a putative truncated tunicamycin resistance gene (*tmrB*, NCBI ref. HBN4657367.1) and a predicted propanediol utilization protein (*pduA*, GenBank ref. QKK72735.1), all flanked by transposable elements. Second, several isolates encoded the β-lactamase gene *bla_TEM-1_*, typically positioned at either the 5’ or 3’ end of the MDR island. Aside from these differences, the larger Ec17R-like regions were essentially the same across all isolates—showing nearly identical nucleotide identity in both coding and intergenic sequences, regardless of collection year.

As expected for the ST131 lineage, and for the P1A strains in particular, all isolates collected from patient P1 in 2019 exhibited ciprofloxacin resistance, probably due to chromosomal mutations in *gyrA* (e.g., *gyrA_D87N_* and *gyrA_S83L_*), *parE*, and *parC* (24). Trimethoprim resistance was most likely associated with the trimethoprim resistant dihydrofolate reductase gene *dfrA17*, which is consistently located near the end of the Ec17R MDR island (**Table 1**, **Figure 2A**) (16). Collectively, genes encoded within Ec17R/Ec17R-like regions are associated with resistance to trimethoprim-sulfamethoxazole, gentamicin, streptomycin, azithromycin, and tetracycline. These MDR islands also contained numerous transposable elements (**Figure 2A**), both internally and at flanking positions, suggesting that Ec17R and some of its components may be mobilized independently of the plasmid backbone. This interpretation aligns with our earlier study, in which Ec17R was detected on both IncF and IncI plasmids (16), consistent with independent horizontal transfer of the entire region. As observed with the PA1 isolates that were collected between 2012 and 2015, less than 5% of those recovered from patient P1 in 2019 were resistant to trimethoprim, suggesting that only a minority of strains within the PA1 population typically carry the Ec17R island. Of note, Ec17R was not detected in any of the 12 urine isolates that were collected in 2019. This may be due to the low number of isolates recovered from urine, reflecting the patient’s asymptomatic status at the time of sample collection.

To further characterize the distribution of Ec17R-like MDR islands, we used the most conserved segment of Ec17R—spanning the cryptic resistance gene *eamA* (25) through *dfrA17* (15,990 bp)—as a query sequence to search the NCBI BLAST core_nt database. This search identified Ec17R and Ec17R-like islands in 496 strains across 18 different Gram-negative species using a cutoff of ≥85% query coverage and ≥99% sequence identity (**Table 1**, **Figure 2B–D**, **Table S1**). *E. coli* was the predominant host; other species included several *Klebsiella* and *Citrobacter* species, *Enterobacter hormaechei, Edwardsiella tarda, Proteus mirabilis, Raoultella ornithinolytica*, and *Vibrio cholerae* (**Figure 2C**). Many of these strains carried ARGs that were exact nucleotide matches to those encoded in Ec17R (**Table 1**). In addition, multiple transposable elements flanking smaller resistance modules exhibited exact or near-exact identity to Ec17R elements. Although smaller constituent resistance islands frequently occur independently of larger MDR regions in publicly available genomes (26–28), the presence of transposable elements bracketing Ec17R/Ec17R-like islands suggests that they may be mobilized as a complete unit. Ec17R-like regions were detected in isolates from 40 countries, with China, South Korea, the United States, Japan, and Switzerland contributing the largest number of strains (**Figure 2B**). Isolation sources encompassed both clinical samples (e.g., blood, feces) and diverse environmental niches (**Figure 2B and D**), underscoring the widespread distribution of Ec17R and Ec17R-like MDR islands.

Within the Ec17R island, two operonic modules that independently confer antibiotic resistance have even more widespread global distributions. The first module was the well-characterized macrolide resistance operon comprising *mph(A), mrx(A),* and *mphR(A)*, which mediates high-level resistance to azithromycin (29, 30) (**Figure S2A, Table S2**). The second module consisted of the following resistance genes: *chrA* (chromate), *sul1* (sulfonamide), *qacE* (quaternary ammonium efflux), *aadA5* (streptomycin), and *dfrA17* (trimethoprim) (**Figure S3A, Table S3**). For both regions, the flanking transposable elements were included in the BLAST query sequence (**Figures S2A and S3A**) and a minimum query coverage and percent identity of ≥99% was used for maximum verisimilitude. The macrolide resistance module was identified in nearly 4000 genomes, excluding reported lab strains and transconjugants, across more than 70 countries and harbored by nearly thirty Gram-negative genera, including *Escherichia, Klebsiella, Morganella, Serratia, Vibrio*, and others (**Figure S2B-D, Table S2**). The larger constituent module was identified in more than 700 genomes across seven Gram-negative genera, including *Escherichia, Klebsiella, Morganella, Proteus,* and several others (**Figure S3B-D, Table S3**). Collectively, these strains originated from essentially all continents except Antarctica (**Figures S2 and S3**). While most isolates had clinical origins—including blood, urine, and feces (**Figure S2B and S3B, Tables S2 and S3**)—the resistance regions were also detected in strains from a wide array of non-clinical sources, including agricultural settings, wildlife, and both freshwater and wastewater.

### Bayesian Inference of Resistance Gene Co-Occurrence in *Escherichia coli*

To investigate whether other patients harbor *E. coli* isolates with globally distributed Ec17R-like islands, we expanded our search to include 267 ExPEC strains that were isolated between 2013 and 2020 from pediatric patients with invasive infections at Primary Children’s Hospital in Salt Lake City, Utah (henceforth referred to as PCH isolates). Most isolates were recovered from blood samples (n = 179), with additional sources including urine, cerebrospinal fluid, respiratory cultures, and others (see methods for exhaustive list). Although the majority of strains belonged to phylogroup B2 (as is typical of ExPEC strains), we observed a diverse representation of phylogroups and sequence types (**Figure 1, Table S4**), as well as heterogeneity in resistance gene content. As all of the PA1 isolates from rUTI patient P1 were clonal and therefore highly similar in both the extent and identity of resistance genes, these strains were excluded from the analysis to avoid bias.

Resistance determinants from the PCH strains were annotated using the ReGAIN pipeline (31), which utilizes NCBI AMRFinderPlus (32, 33); from these predictions, a presence/absence matrix of genes across the 267 PCH genomes was generated. To reduce noise, very high (>260) and very low (<5) prevalence genes were excluded. Of note, two strains were identified as harboring the colistin (polymyxin E) resistance determinants *mcr-1* or *pmrB_V161G_* (**Table S5**), but these were not considered in our analysis due to their low prevalence. Hierarchical clustering of these data (Jaccard similarity, **Figure 3A and B**) revealed several clear patterns of ARG co-occurrence. Principal coordinate analysis (PCoA, Jaccard similarity; **Figure 3C**) further highlighted intrinsic patterns of gene clustering, including variant-specific associations. For example, the trimethoprim resistance gene *dfrA17* and the streptomycin resistance gene *aadA5* preferentially clustered together, distinct from *dfrA12* and *aadA2*, which formed a separate cluster. Similarly, the tetracycline efflux transporter *tet(A)* clustered with *dfrA17* and *aadA5*, whereas *tet(B)* clustered with *dfrA5* and *aadA1*. Notably, only *dfrA1* and *dfrA17* consistently clustered with sulfonamide resistance genes, despite trimethoprim-sulfamethoxazole being used clinically as a combinatorial therapy. Several clusters also contained genes conferring resistance to multiple antibiotic classes, consistent with localization patterns within MDR islands (**Figure 3C**). While most heavy metal resistance genes clustered independently, some—including *merR, merP, merT, merC, terZ, terD, terW,* and *arsD*—were found in proximity to ARGs (**Figure 3C**). Alarmingly, ten PCH isolates carried large MDR islands that closely resembled Ec17R, including two isolates in which all resistance genes were identical to those within Ec17R at the nucleotide level (**Figure S4)**.

**Figure 3.**
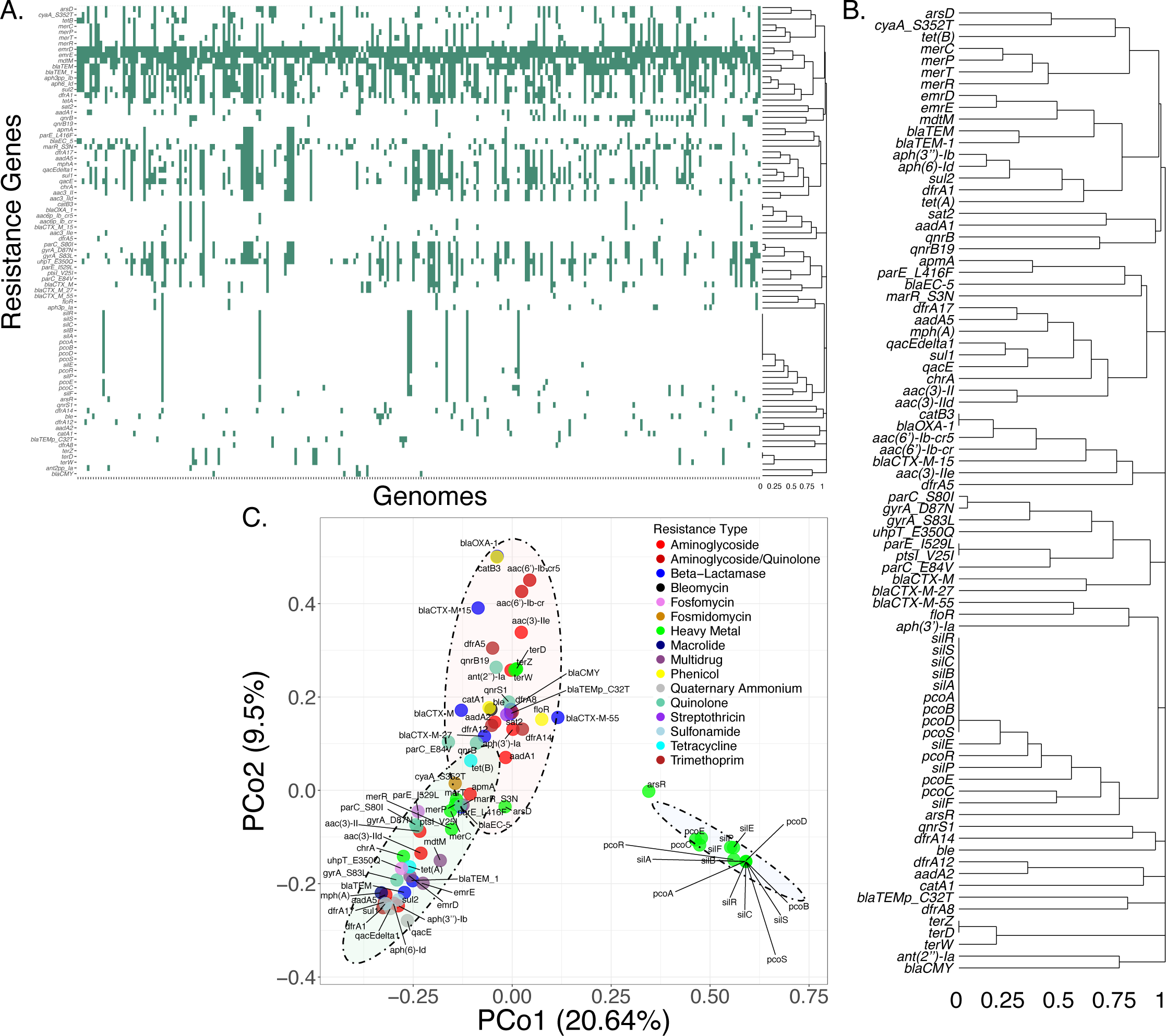
Patterns of resistance gene co-occurrence in *E. coli* strains isolated from hospitalized pediatric patients. (**A**) Heatmap and dendrogram (right) generated using hierarchical clustering (Jaccard) of ARG presence/absence correlations across genomes from 267 sequenced PCH isolates. (**B**) Detail of the dendrogram from panel A. (**C**) PCoA (Jaccard) plot showing clustering of ARGs and HMRGs, with genes colored by resistance class. Ellipses represent 95% confidence intervals.

Although clustering approaches provide insight into similarity and correlation, they are limited in their ability to capture conditional dependencies. To more rigorously quantify co-occurrence, we applied a Bayesian network structure learning model. Bayesian networks are probabilistic graphical models that compute relationships between variables based on conditional dependencies (34, 35). Using our PCH dataset, we constructed a Bayesian network using a modified ReGAIN pipeline and derived pairwise conditional probabilities, absolute risk differences, and relative risk ratios for each gene pair (**Figure 4, Table S6**). Conditional probability was defined as the probability that Gene B is present given Gene A, P(A|B). Absolute risk difference was defined as the probability that Gene B is present given Gene A, P(A|B) minus the conditional probability that Gene B is absent given Gene A, P(A|¬B) (i.e., the conditional probability of co-occurrence minus the conditional probability of single occurrence). Relative risk was defined as the ratio P(A|B) / P(A|¬B), where values >1 indicate positive relationships, <1 indicate negative relationships, and =1 indicate neutral relationships.

**Figure 4.**
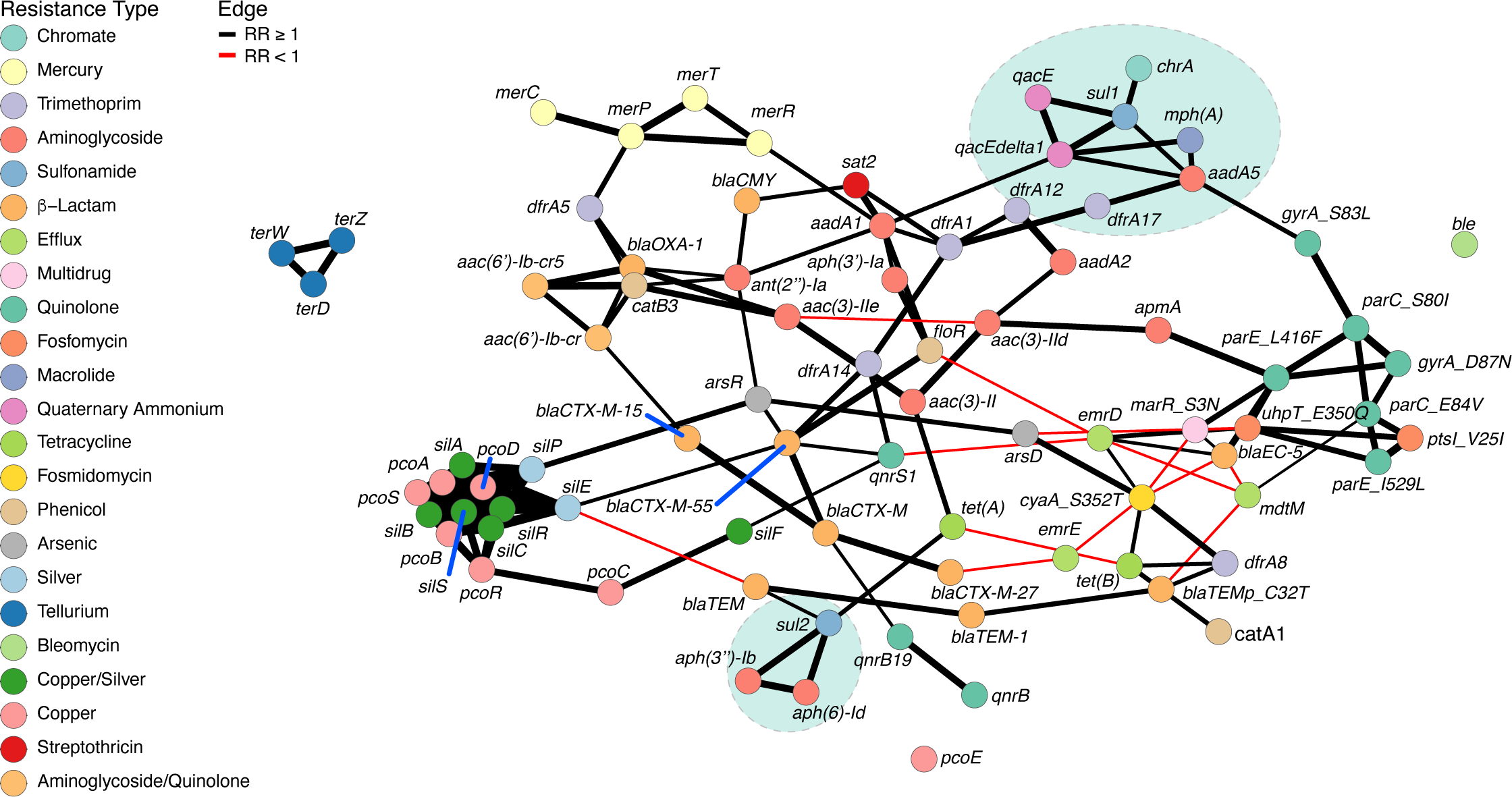
Bayesian network of resistance genes identified in PCH *E. coli* isolates reveals the co-occurrence of multiple antibiotic and heavy metal resistance genes. Nodes are colored by resistance type; edge colors reflect either positive (black) or negative (red) probabilistic relationships based on relative risk ratios. Edge width reflects absolute risk confidence intervals, where wider edges indicate higher confidence. *pcoE* and *ble* represent isolated nodes. Relative risk >1 indicates positive relationship; <1 indicates negative relationship; =1 indicates neutral relationship. Green circles highlight resistance genes associated with Ec17R.

To validate our approach, we first examined genes known to co-occur, including *aph(3’’)-IIb, aph*(*6*)*-Id,* and *sul2* (36–38). As expected, we observed strong positive relationships: *aph(3’’)-IIb* with *aph*(*6*)*-Id* (abs. risk diff. 0.89; rel. risk 26.33), *aph(3’’)-IIb* with *sul2* (abs. risk diff. 0.79; rel. risk 7.88), and *aph*(*6*)*-Id* with *sul2* (abs. risk diff. 0.78; rel. risk 7.84). Within the network, these three genes form a tight cluster indicative of strongly associated variables (**Figure 4**). We also identified unexpected associations, such as between the chloramphenicol resistance gene *catB3* and the cephalosporinase *bla_OXA-1_* (abs. risk diff. 0.94; rel. risk 989.24), and between the β-lactamase *bla_CTX-M-55_* and the florfenicol resistance gene *floR* (abs. risk diff. 0.75; rel. risk 85.44). Intriguingly, a recent report describes ST131 isolates from Armenia that encode an MDR island carrying *bla_CTX-M-15_, aac*(*3*)*-IIe, catB3,* and *aac(6’)-Ib-cr6* (39). In the PCH dataset, we observed positive relationships among the same genes, though the quinolone resistance determinant *aac(6’)-Ib-cr6* was represented by *aac(6’)-Ib-cr5* in the PCH isolates, potentially reflecting regional or temporal variation among strains (40–43). These findings indicate that co-occurrence of chloramphenicol and cephalosporin resistance genes like *bla_OXA-1_* and *bla_CTX-M-15_* is not unique to U.S. isolates, as suggested by previous studies (44). Importantly, many Ec17R constituents form tight clusters within the network, reinforcing their empirical co-occurrence architecture (**Figure 4**).

Finally, we investigated co-occurrence between antibiotic and heavy metal resistance determinants. Positive relationships included a suite of mercury resistance genes (*merC, merP,* and *merT*) with the trimethoprim resistance gene *dfrA5* (abs. risk diff. 0.45, 0.49, 0.45; rel. risk 4.02, 5.00, 3.56, respectively) and the arsenic resistance gene *arsD* with the fosfomycin resistant adenylate cyclase gene *cyaA_S352T_* (abs. risk diff. 0.66; rel. risk 7.88) and *tet(B)* (abs. risk diff. 0.18; rel. risk 2.51). Of particular note, the chromate resistance gene *chrA* exhibited elevated co-occurrence with ARGs across multiple classes, including aminoglycosides (gentamicin, streptomycin) and macrolides. However, in general, heavy metal resistance genes displayed few strong probabilistic relationships with ARGs (**Table S6**). Indeed, the most common heavy metal resistance genes (*pco* and *sil* family genes) formed the tightest associations with other heavy metal resistance genes (**Figure 4**), with the exception of *pcoE*, which represents an isolated node in the network. Notably, transcriptional regulators associated with heavy metal resistance (*merR, arsR*) did exhibit strong relationships with certain ARGs (Figure 4, Table S6).

### Survival of rUTI *Escherichia coli* Isolates under β**-**Lactam Exposure

Given the long-term maintenance of the clonal PA1 population within patient P1 despite extensive treatments with antibiotics for which the PA1 strains encode no known resistance mechanisms (16), we reasoned that these pathogens may be intrinsically recalcitrant to some antibiotics. To address this possibility, antibiotic survival assays were performed as previously described (21, 23) using four of the PA1 isolates along with the well-studied reference ExPEC strain CFT073 (21, 45). Of the four PA1 isolates tested, two harbored either Ec17R (U15A) or a highly similar Ec17R-like island (U19fE, missing *aac*(*3*)*-IId* module), and two lacked any version of the MDR island (U19fL, U19fI). All of these strains were genetically and phenotypically sensitive to the β-lactam antibiotics ampicillin and ertapenem (**Table S7**).

For our survival assays, CFT073 and each of the PA1 strains were challenged in broth culture for 5 hours with either ampicillin or ertapenem present at 10–30x their minimum inhibitory concentrations (MICs) (**Table S7**). These assays provide a measurement of antibiotic recalcitrance, which encompasses both antibiotic persistence and tolerance phenotypes (46). In the presence of ampicillin, strain U15A survived at markedly higher levels than the other patient isolates, including U19tE, which has a nearly identical Ec17R island (**Figure 5A**). In assays with ertapenem, survival levels varied moderately among the patient isolates, and also did not appear to depend on the presence of Ec17R (**Figure 5B**). All of the PA1 isolates survived at much higher levels than the CFT073 reference strain (**Figure 5**). Importantly, in the absence of any antibiotics, U15A, U19fE, U19fL, and U19fI all grew at similar rates, and all were faster and reached greater densities than CFT073 (**Figure S5**). These findings suggest that slower growth does not necessarily promote ExPEC survival in these assays, though reduced growth rates are thought to render some bacteria less susceptible to β-lactams and other cell wall-targeting antibiotics (47). Instead, our results point to as-yet unidentified, Ec17R-independent mechanism(s) that enable survival of the PA1 isolates during exposure to antibiotics for which they are sensitive.

**Figure 5.**
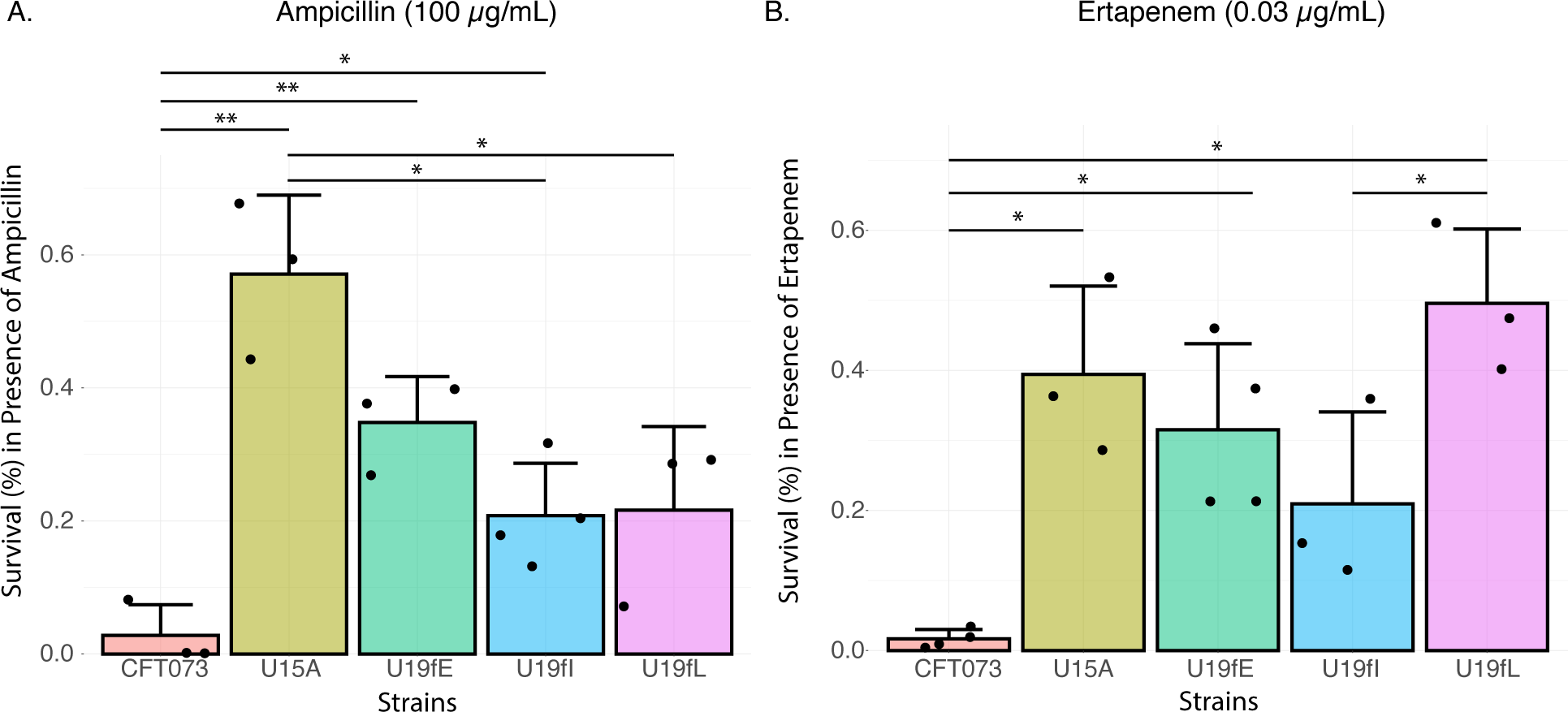
PAI isolates are able to survive high levels of ampicillin and ertapenem despite lacking known resistance mechanisms for these antibiotics. (**A**) Percent survival in the presence of ampicillin. (**B**) Percent survival in the presence of the ertapenem. Strains U15A, U19fE, U19fI, and U19fL do not encode any class A or D β-lactamase genes but survive at much higher levels than reference strain CFT073. 0.6% survival equates to 7.5×10^6^ CFU. *, p < 0.05, **, p < 0.01. Significance was calculated by Student’s *t*-tests using 3−4 biological replicates.

To identify potential genetic factors that can impact the recalcitrance of the PA1 strains to antibiotics, we leveraged the high genetic homogeneity of these isolates (**Figure 1B**) (16) to perform pairwise genomic comparisons. Specifically, we compared gene content of the two strains harboring Ec17R/Ec17R-like regions (U15A and U19fE), which differed in their survival rates in the presence of ampicillin, and the two Ec17R-negative isolates, U19fL and U19cL, that differed greatly in survival assays with ertapenem (**Figure 5**). Our analysis revealed that U15A has 19 predicted genes that were distinct from those in U19fE (**Table S8**). These included multiple hypothetical genes, the aminoglycoside resistance operon containing *aac*(*3*)*-IId,* and the putative truncated tunicamycin resistance gene *tmrB*, as well as a few genes associated with secretion and horizontal gene transfer (*traI*, *traE*, *traF,* and *traH*). Eight genes were found to be uniquely encoded by U19fL, relative to U19fI (**Table S9**). One of these encoded a leader peptide (IvbL) that reportedly helps control expression of genes involved in isoleucine and valine biosynthesis, while the remaining U19fL-associated genes were annotated as hypotheticals. Though several of the genes identified by this comparative analysis have the potential to impact the bacterial envelope and metabolic pathways, the means by which these factors might affect antibiotic recalcitrance within the PA1 lineage and beyond will require additional investigation. Altogether, our results stress the need to consider both antibiotic resistance and recalcitrance mechanisms when considering MDR epidemiology.

## DISCUSSION

Multidrug resistance is not a localized issue but a global crisis, affecting healthcare systems and populations worldwide (1, 5, 8, 48). Here, we applied robust statistical tools to map patterns of resistance gene co-occurrence, providing predictive value for understanding how multidrug resistance emerges and spreads. Applying a novel Bayesian network structure learning approach to analyze ARG co-occurrence in clinical *E. coli* isolates enabled us to quantify conditional dependencies and relative risks between gene pairs. This approach uncovered both expected associations—such as among aminoglycoside–sulfonamide resistance genes—and striking, unexpected relationships like those between chloramphenicol and cephalosporin resistance determinants. The latter is particularly concerning, as chloramphenicol is rarely used clinically in the U.S. (49), suggesting that global strain movement via, for example, international travel or foodborne dissemination (38, 50) may underlie these patterns. Such findings underscore the utility of probabilistic frameworks for monitoring resistance gene relationships across populations, while also highlighting limitations such as rare or ubiquitous determinants that can introduce noise, requiring careful curation or complementary analyses. This is exemplified by the detection of polymyxin resistance determinants (*mcr-1, pmrB_V161G_*) in our PCH pediatric isolates, which is especially worrisome given the reliance on polymyxin as a drug of last-resort (51, 52).

Beyond assessing ARG co-occurrence, our analysis extended to HMRGs. MDR islands may cluster both ARGs and HMRGs, enabling co-selection under diverse environmental pressures. We identified several instances of ARG and HMRG co-occurrence, most notably between the chromate resistance gene *chrA*, which is a constituent of the Ec17R island (**Figure 2A**), and multiple ARGs. These findings are consistent with prior reports in *Salmonella* (20, 53, 54). While most HMRGs in our dataset did not exhibit strong relationships with ARGs, our results reinforce the importance of environmental selective pressures—such as heavy metal pollutants—in shaping resistance architectures. Cataloging and prioritizing such associations will help identify candidate genes for future experimental studies and may inform surveillance strategies by linking environmental and clinical reservoirs.

Critically, ARGs represent only one facet of bacterial survival in the face of antibiotics. Our *in vitro* survival assays revealed that different PA1 isolates tolerated exposure to ampicillin and ertapenem at concentrations well above their MICs, despite lacking classical resistance determinants (see **Figure 5**). The survival phenotypes of the PA1 isolates did not correlate with slower growth kinetics, which has been associated with elevated antibiotic recalcitrance in some bacteria (47). Comparative genomics of PA1 isolates that vary in their antibiotic survival phenotypes identified candidate genes potentially linked to antibiotic recalcitrance, including secretion system components and multiple hypothetical proteins. Though several of these genes have the potential to directly or indirectly impact the bacterial envelope, metabolic pathways, or stress responses, the mechanisms by which these factors might affect antibiotic recalcitrance within the PA1 lineage and beyond requires additional investigation.

Clinically, antibiotic recalcitrance likely contributes to cycles of rUTIs, as experienced by patient P1. In her case, ertapenem treatment provided a fairly long reprieve from UTIs caused by resident PA1 strains, but surviving sub-populations eventually re-established symptomatic infections (16). This sort of confounding situation, in which antibiotics provide only fleeting relief, is not uncommon among individuals who suffer from rUTIs (55), and likely often involves the resurgence of recalcitrant sub-populations that are only transiently suppressed by antibiotic treatments (16). Our results from survival assays with the PA1 strains further emphasize that antibiotic recalcitrance (including both tolerance and persistence mechanisms) likely contributes to treatment failures and may be as clinically significant as resistance, yet recalcitrance is not captured by standard susceptibility testing.

Taken together, our study advances three key insights. First, Bayesian networks provide a powerful lens for mapping probabilistic relationships among resistance genes, revealing both canonical and unexpected associations. Second, co-occurrence between ARGs and some HMRGs highlights the potential for environmental co-selection of resistance determinants. Third, antibiotic survival in the absence of resistance underscores the urgent need to better integrate tolerance and persistence (recalcitrance) into our framework for understanding treatment failure and disease recurrence. As genomic datasets expand, applying probabilistic modeling to well-annotated strain collections will allow increasingly confident mapping of co-occurrence patterns. Ultimately, coupling such approaches with mechanistic studies of antibiotic recalcitrance will better equip us to anticipate the trajectories of multidrug resistance and to refine stewardship and diagnostic strategies.

## MATERIALS AND METHODS

### Strain Acquisition and Isolation

From patients who were hospitalized at Primary Children’s Hospital (PCH) in Salt Lake City, Utah between 2013 and 2020, 267 *E. coli* isolates were collected from blood (n=179), urine (n=15), cerebral spinal fluid (n=12), peritoneal fluid (n=12), abscess (n=11), respiratory culture (n=12), feces (n=7), sputum (n=3), pus (n=3), wound (n=3), fluid culture (n=1), pleural fluid (n=2), tissue culture (n=3), bile (n=1), bone culture (n=1), kidney (n=1), and perihepatic fluid (n=1).

In 2019, 24 fecal and 12 urine isolates were collected from rUTI patient P1 at the University of Utah Urology Clinic. Samples were plated on MacConkey agar ± ciprofloxacin (10 μg/mL) or trimethoprim (20 μg/mL) as previously described (16). The patient had not received any antibiotics for approximately three weeks prior to sample collection and was asymptomatic for UTI. The 12 urine isolates were recovered only after concentrating the patient’s urine sample via brief centrifugation prior to plating. All bacterial isolates were stored at -80°C using Viabank^™^ bacterial storage beads.

### Genome Preparation and Assembly

Genomic DNA was harvested from each bacterial isolate using a Qiagen DNeasy^®^ Blood & Tissue Kit. Short-read sequences were obtained using an Illumina NovaSEQ6000 instrument through the University of Colorado Anschutz Genomics and Microarray Core. Sequencing parameters were 151 bp paired-end reads at a depth of 30x. Adapter sequences and PhiX were removed from all reads with BBDuk, which is part of the Bbtools suite (56). Reads were trimmed using Qtrim and assessed for quality using FastQC (57, 58). *De novo* sequence assembly was performed using SPAdes (59), and assemblies were annotated using Prokka (60). Long-read sequencing and genome assembly of *E. coli* isolates U15A, U19fE, U19fI, and U19fL was performed using Oxford Nanopore Technology by Plasmidsaurus (Eugene, Oregon).

### Genomic Analyses of Ec17R Island

The most conserved 15,990 base pair region in Ec17R was used as a nucleotide query to the NCBI BLAST core_nt database first against *E. coli* specifically, yielding 414 genomes fitting a minimum cutoff of ≥85% query coverage and ≥99% sequence identity. Next, the same query was repeated, excluding *E. coli* from the search. This returned 82 genomes meeting our inclusion criteria. Genome metadata was extracted from GenBank files using seqforge v1.0.4 (61). Figures were constructed in R using the following packages: ggplot2 v3.4.2, ggalluvial v0.12.5 and forcats v1.0.0. Smaller genomic operons were analyzed in similar fashion, except that both the query coverage and sequence identity cutoffs were each set to ≥99%. Core gene alignments were generated using Panaroo (v1.5.2) (62), and then used to create maximum likelihood phylogenetic trees via IQ-TREE (v3.0.1) (63) using the GTR+G model of nucleotide substitution and 1000 bootstraps. Manual annotation of resistance genes in the Ec17R island was performed using NCBI BLAST+ (64), as well as the NCBI BLAST web application. Comparative genomics was performed using Anvi’o (v8) (65). Unique genes were extracted using a custom Bash pipeline and annotated using the NCBI BLAST protein database. Serotypes were predicted using ECTyper (v1.0.0rc1) (66). Multi-locus sequence typing of strains were performed using mlst (v2.23.0) (https://github.com/tseemann/mlst) (67).

### Resistance Gene Identification, Gene-Gene Correlations, and Statistical Analyses

Antibiotic resistance genes were identified using the Resistance Gene Association and Inference Network (ReGAIN) bioinformatic pipeline (31), which utilizes NCBI AMRfinderPlus (v4.0.23) (33) using the *Escherichia* organism flag, with the exception of the chromate resistance gene, *chrA*, which was identified within the genomic population using the ReGAIN Curate pipeline (https://github.com/ERBringHorvath/regain_CLI). The ChrA protein sequence identified in *E. coli* strain U15A, plasmid pU15A_A (NZ_CP035721.1) was used as a query sequence. Statistical analyses were performed in Rstudio using the following packages: vegan (v2.6-8) (68), bnlearn (v 5.0.1) (34), gRain (v1.4.5) (69), gRbase (v2.0.3) (70), stats (v4.2.1), and ggpubr (v0.6.0). Results were visualized using ggplot2 (v3.4.2) and ggbreak (v.0.1.1) (71). Hierarchical clustering and principal coordinate analyses were performed using the Jaccard similarity measure. Discrete Bayesian network structure learning was performed using bnlearn and gRain using 1000 bootstraps and 1000 resamples with replacement. To reduce noise in the Bayesian network, genes occurring less than 5 times or more than 260 times in the PCH bacterial genomes were not considered during network learning. The cephalosporinase, *bla_EC_,* was identified in each of the 267 PCH isolates; because discrete binary Bayesian network structure learning requires all variables to be both ‘present’ and ‘absent’ (i.e., variables cannot be ubiquitous), *bla_EC_* was removed from the dataset prior to structure learning.

### Minimum Inhibitory Concentrations and Growth Kinetics

Minimum inhibitory concentrations (MIC) were quantified using Liofilchem MIC Test Strips (Fisher Scientific). Briefly, *E. coli* strains CFT073, U15A, U19fE, U19fI, and U19fL were grown in triplicate from frozen stocks overnight in Lysogeny Broth (LB) (37°C, 220 rpm), and then back diluted and normalized to a concentration of 5×10^6^ CFU/mL. Aliquots (100 µL) of each culture were then spread on LB agar and MIC test strips were applied. After an 18-hour incubation at 37°C, MIC values were read using the integrated scale on each test strip. Preparation of *E. coli* strains for growth kinetics followed the same initial protocol as with MIC studies. After back dilution and normalization, *E. coli* strains CFT073, U15A, U19fE, U19fI, and U19fL were plated using biological triplicates on a 96-well plate. Plates were incubated at 37°C and OD_600_ was measured every 30 minutes for 24 hours, with brief shaking occurring immediately before measurement.

### Antibiotic Survival Assays

Survival assays were carried out essentially as described previously (21). In short, overnight cultures of each bacterial isolate were grown from frozen stocks shaking at 220 rpm at 37°C and then back diluted to an OD_600_ of 1.0. Next, 100 µL of each OD-normalized culture was added to 4.9 mL of fresh LB ± ampicillin (100 µg/mL) or ertapenem (0.03 mg/L). After a 5-hour incubation period (220 rpm, 37°C), 1 mL of each culture was extracted and pelleted by centrifugation at 16,000 x *g* for 1 minute; cell pellets were washed in cold Phosphate Buffered Saline (PBS, pH 7.4) and enumerated by plating serial dilutions onto LB agar. The numbers of bacteria that survived the antibiotic treatments were calculated as percentages of the bacteria recovered from untreated control cultures. Statistical tests were carried out in R using Student’s *t*-tests with a Benjamini-Hochberg post-hoc analysis. *P* values of <0.05 were considered significant.

## Supporting information

Table S1: Strains containing Ec17R-like MDR islands, with associated information on sources, regions of isolation, host organisms, and collection date

Table S2: Strains containing a Streptomycin/Sulfonamide operon as seen in Ec17R, with associated information on sources, regions of isolation, host or

Table S3: Strains containing a Macrolide resistance operon as seen in Ec17R, with associated information on sources, regions of isolation, host organi

Table S4: Multi-locus sequence type, phylogroup, H-type, and O-type of pediatric patient E. coli isolates

Table S6: Probability metrics of resistance gene co-occurrence in PCH E. coli population

## ACKNOWLEDGEMENTS and FUNDING SOURCES

We thank Qin Zhou and Shannon Nielsen for assistance with processing the clinical samples. This work was supported by a University of Utah 3i graduate research fellowship to ERBH, NIH Genetics T32 training grant GM007464 to OJM, University of Utah Research Foundation funds to JMW, a Margolis Foundation grant to WJB, MAM, and JMW, by the Department of Defense award W81XWH-22-1-0800 (SC210103) to WJB, SML, JMW and MAM, and by NIH grant GM134331 to MAM.

## Ethics Approval Statement

The research described in this study was undertaken with approval from the University of Utah Institutional Biosafety Committee and the Institutional Review Board (IRB_00097637, IRB_00070563).

## Data Availability Statement

All genomes have been made available under BioProject accession number 000000000 (TBD).

## AUTHOR CONTRIBUTIONS

**Elijah R. Bring Horvath:** conceptualization (equal); bioinformatics (lead); survival assays (lead); data analysis (lead); writing – original draft (lead); writing – review and editing (equal). **O’Connor Matthews:** bacterial culture and preparation (lead); DNA isolation (lead); bioinformatics (equal); writing – review and editing (equal). **William J. Brazelton**: conceptualization (equal); bioinformatics (equal); sequencing and genome assembly (equal); data analysis (equal); writing – review and editing (equal). **Edgar Javier Hernandez:** conceptualization (equal); bioinformatics (equal); writing – review and editing (equal). **Hillary Crandall:** strain collection and isolation (equal). **Sara M. Lenherr:** strain collection and isolation (equal). **Jaclyn M. Winter**: conceptualization (equal); project administration (lead); data analysis (equal); writing − original draft (equal); writing − review and editing (equal). **Matthew A. Mulvey:** conceptualization (equal); project administration (lead); data analysis (equal); writing − original draft (equal); writing − review and editing (lead).

