## Supplemental Data 1 for "Global Footprint of the Multidrug Resistance Island Ec17R and Resistance Gene Co-Occurrence in Pathogenic *Escherichia coli* Isolates"

### Table of Contents

#### Supplementary Figures


Figure S1: Multidrug resistance island identified in rUTI patient *E. coli* isolates S3
Figure S2. Distribution of Ec17R-constituent macrolide resistance operon S4
Figure S3: Distribution of Ec17R-constituent multidrug resistance region S5
Figure S4: PCH isolates harboring Ec17R or Ec17R-like MDR regions S6
Figure S5: Growth curves of *E. coli* strains CFT073, U15A, U19tE, U19cL, and U19cI in LB broth S11


#### Supplementary Tables


Table S1: Strains containing Ec17R-like MDR islands, with associated information on sources, regions of isolation, host organisms, and collection dates, if known S7
Table S2: Strains containing a Streptomycin/Sulfonamide operon as seen in Ec17R, with associated information on sources, regions of isolation, host organisms, and collection dates S7
Table S3: Strains containing a Macrolide resistance operon as seen in Ec17R, with associated information on sources, regions of isolation, host organisms, and collection dates S7
Table S4: Multi-locus sequence type, phylogroup, H-type, and O-type of pediatric patient *E. coli* isolates S7
Table S5: PCH (pediatric) isolates encoding genes associated with polymyxin resistance S8
Table S6: Probability metrics of resistance gene co-occurrence in PCH *E. coli* population S9
Table S7: Minimum inhibitory concentrations of ampicillin and ertapenem against rUTI patient-derived P1A *E. coli* isolates S10
Table S8: Genes encoded by strain U15A that are not encoded by strain U19fE S12
Table S9: Genes encoded by strain U19fL that are not encoded by strain U19fI S13

**Supplemental Reference** S14

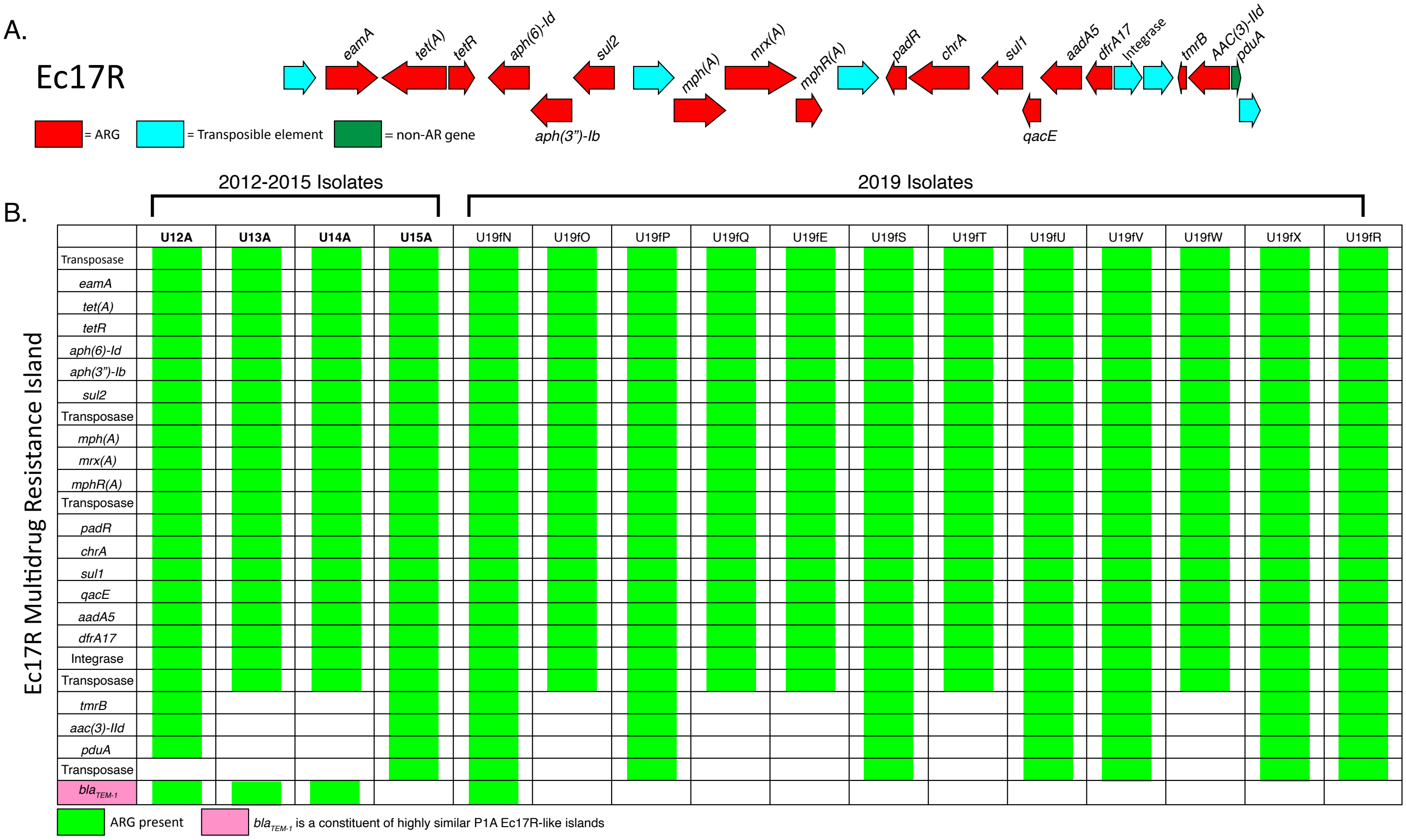


**Figure S1.** Features of the Ec17R MDR island carried by the PA1 isolates. (**A**) MDR region identified in strain U15A (1). (**B**) Comparison of MDR regions between strains isolated from the rUTI patient. All Trimethoprim-resistant strains that were sequenced from patient P1 carried the Ec17R island. *bla*$_{TEM-1}$ is found in some P1A Ec17R-like islands, but not in Ec17R itself.

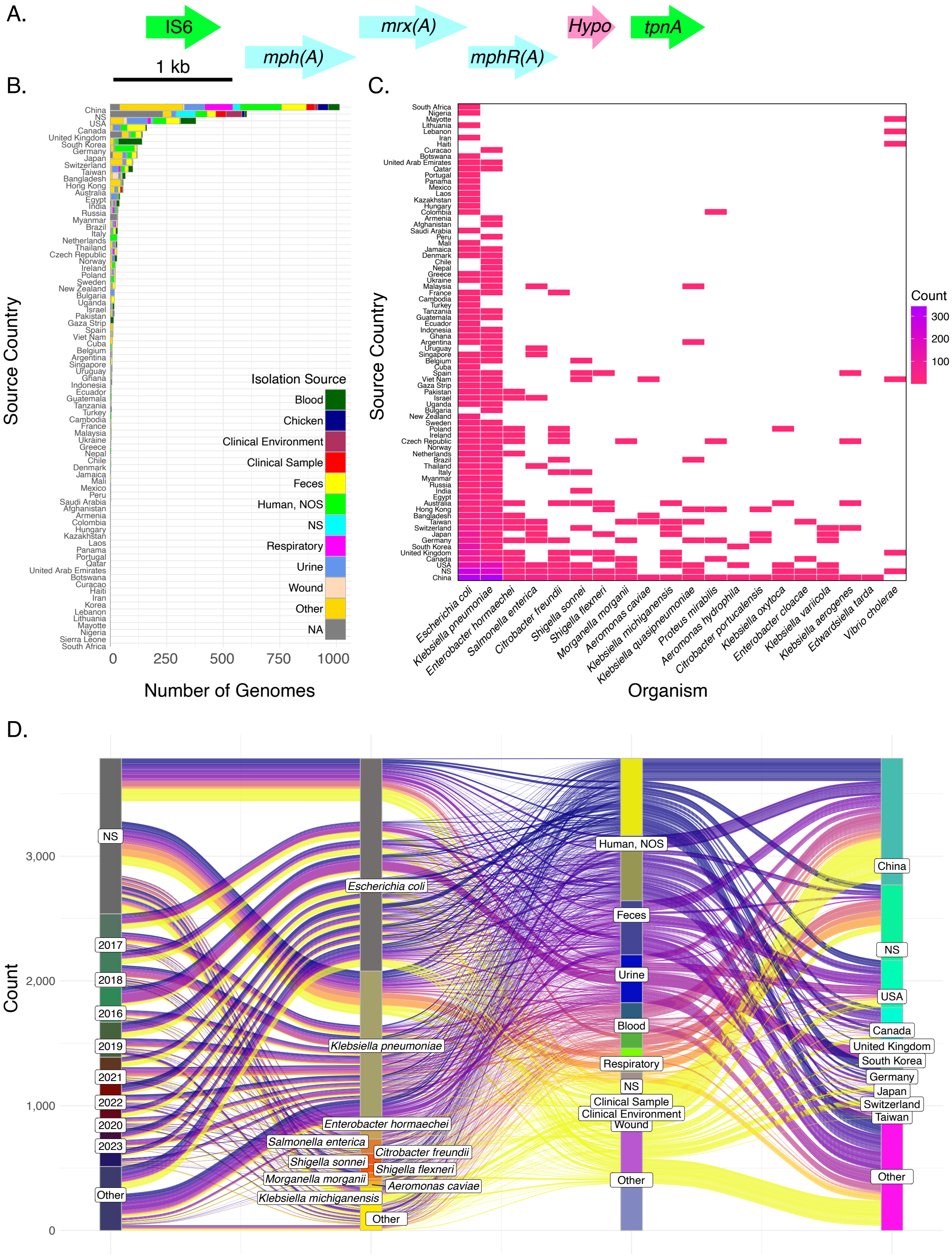


**Figure S2.** Global footprint of Ec17R constituent macrolide resistance operon**.** (**A**) Macrolide resistance operon, including transposable elements, identified in *E. coli* strain U15A (plasmid pU15A_A) isolated in Salt Lake City, Utah from a recurrent UTI patient. Arrows represent open reading frames and point in the direction of transcription. Blue: predicted resistance genes. Green: transposable elements. Fuchsia: hypothetical. (**B-D**) Sources of strains harboring macrolide resistance islands identified via NCBI nucleotide blast. (**B**) Ten most common isolation sources and country. (**C**) Species distribution across source countries. (**D**) Top ten collection years, organisms, isolation sources, and source countries. NS, not specified; NOS, Not Otherwise Specified.

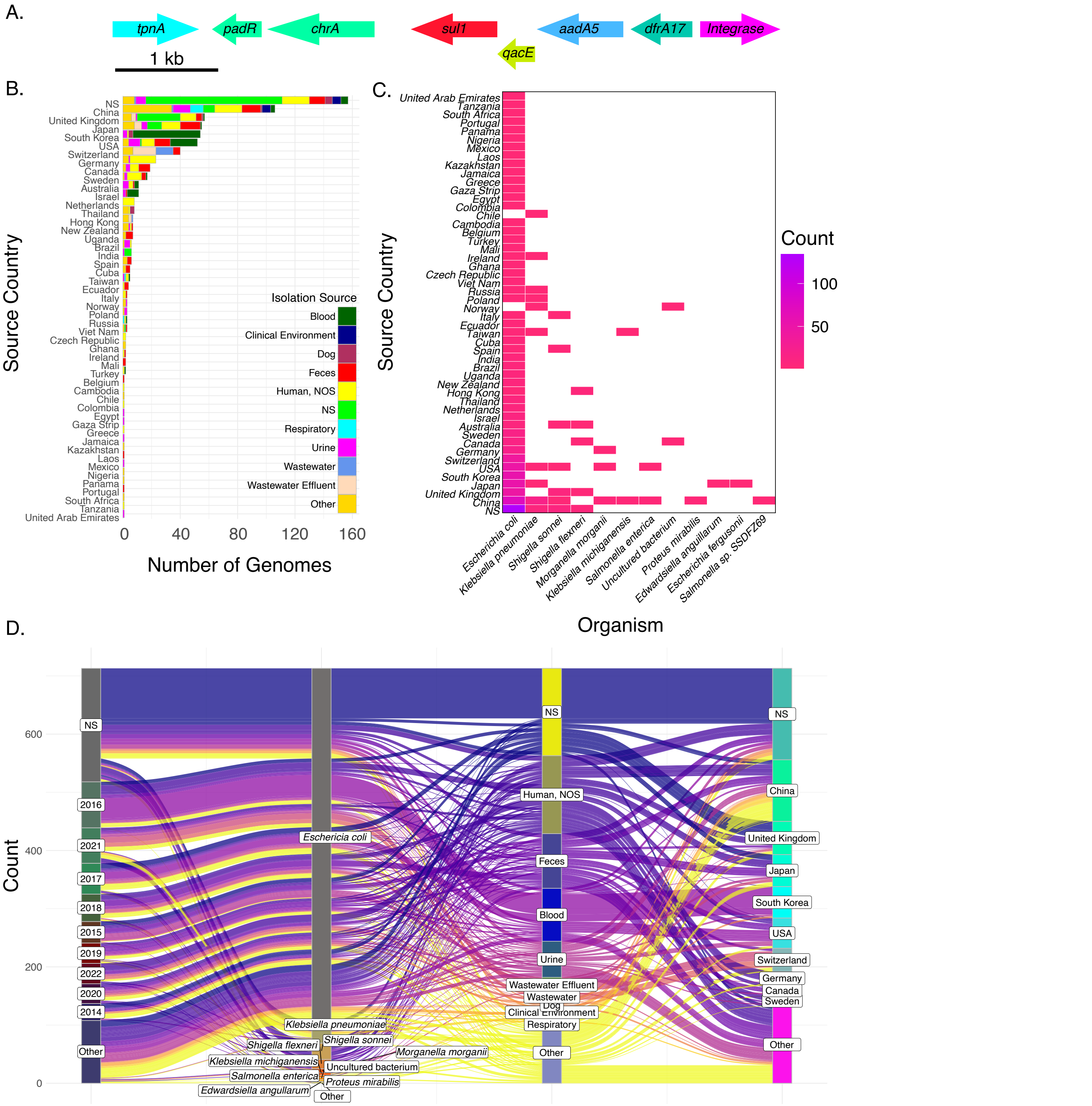


**Figure S3.** Global footprint of Ec17R constituent multidrug resistance region. (**A**) Resistance region, including transposable elements, identified in *E. coli* strain U15A (plasmid pU15A_A) isolated in Salt Lake City, Utah from a recurrent UTI patient. Arrows represent open reading frames and point in the direction of transcription. Aquamarine: chromate resistance. Red: sulfonamide resistance. Chartreuse: quaternary ammonium efflux. Blue: streptomycin resistance. Sea green: trimethoprim resistance. Cyan: IS6-family transposase. Violet: integrase. (**B-D**) Sources of strains harboring resistance islands identified via NCBI nucleotide blast. (**B**) Ten most common isolation sources and country. (**C**) Species distribution across source countries. (**D**) Top ten collection years, organisms, isolation sources, and source countries. NS, not specified; NOS, Not Otherwise Specified.

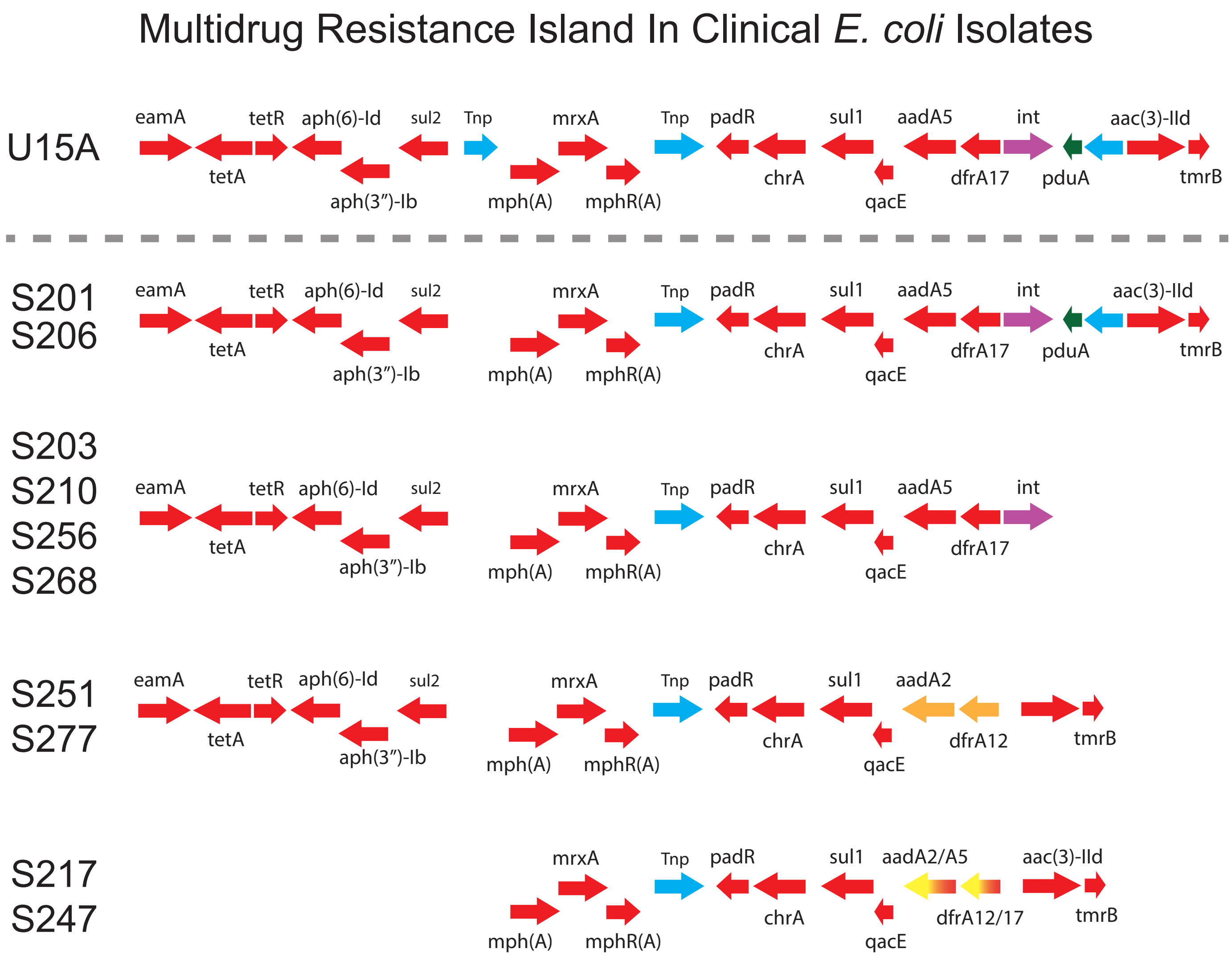


**Figure S4**. Primary Children's Hospital (PCH) isolates harboring Ec17R or Ec17R-like MDR regions. Strain names are indicated on the left. Top shows the Ec17R island from UTI isolate U15A as a reference. Multi-colored arrows (yellow and red) indicate genes that could be one of two variants.

**Table S1:** Strains containing Ec17R-like MDR islands, with associated information on sources, regions of isolation, host organisms, and collection dates, if known.
**Table S2:** Strains containing a Streptomycin/Sulfonamide operon as seen in Ec17R, with associated information on sources, regions of isolation, host organisms, and collection dates.
**Table S3:** Strains containing a Macrolide resistance operon as seen in Ec17R, with associated information on sources, regions of isolation, host organisms, and collection dates.
**Table S4:** Multi-locus sequence type, phylogroup, H-type, and O-type of pediatric patient *E. coli* isolates

Due to the size of these tables, they are available as separate downloadable spreadsheets.

https://figshare.com/articles/dataset/Global_Distribution_of_Ec17R_Multidrug_Resistance_Islands_and_Probabilistic_Mapping_of_Resistance_Gene_Co-Occurrence_in_Clinical_i_Escherichia_coli_i_Supplemental_Tables/30268069

**Table S5.** PCH (pediatric) isolates encoding genes associated with polymyxin resistance.

| Strain | Gene | NCBI RefSeq | % Identity |
|---|---|---|---|
| S305 (*E. coli*) | *mcr-1* | WP_034169413.1 | 99 |
| S221 (*E. coli*) | *pmrB_V161G* | WP_001300761.1 | 100 |

**Table S6.** Probability metrics of resistance gene co-occurrence in PCH *E. coli* population

Due to the size of this table, it is available as a separate downloadable spreadsheet.

https://figshare.com/articles/dataset/Global_Distribution_of_Ec17R_Multidrug_Resistance_Islands_and_Probabilistic_Mapping_of_Resistance_Gene_Co-Occurrence_in_Clinical_i_Escherichia_coli_i_Supplemental_Tables/30268069

**Table S7.** Minimum inhibitory concentrations of ampicillin and ertapenem against rUTI patient-derived P1A *E. coli* isolates.

| Strain | MIC Ertapenem (μg/mL) | SD | MIC Ampicillin (μg/mL) | SD |
|---|---|---|---|---|
| CFT073* | 0.004 | ± 0.0003 | 2.67 | ± 0.58 |
| U15A | 0.005 | ± 0.003 | 4.67 | ± 1.15 |
| U19tE | 0.006 | ± 0.002 | 3.33 | ± 0.58 |
| U19cI | 0.006 | ± 0.0006 | 3.50 | ± 0.50 |
| U19cL | 0.009 | ± 0.0003 | 4.03 | ± 0.058 |

*Reference ExPEC strain

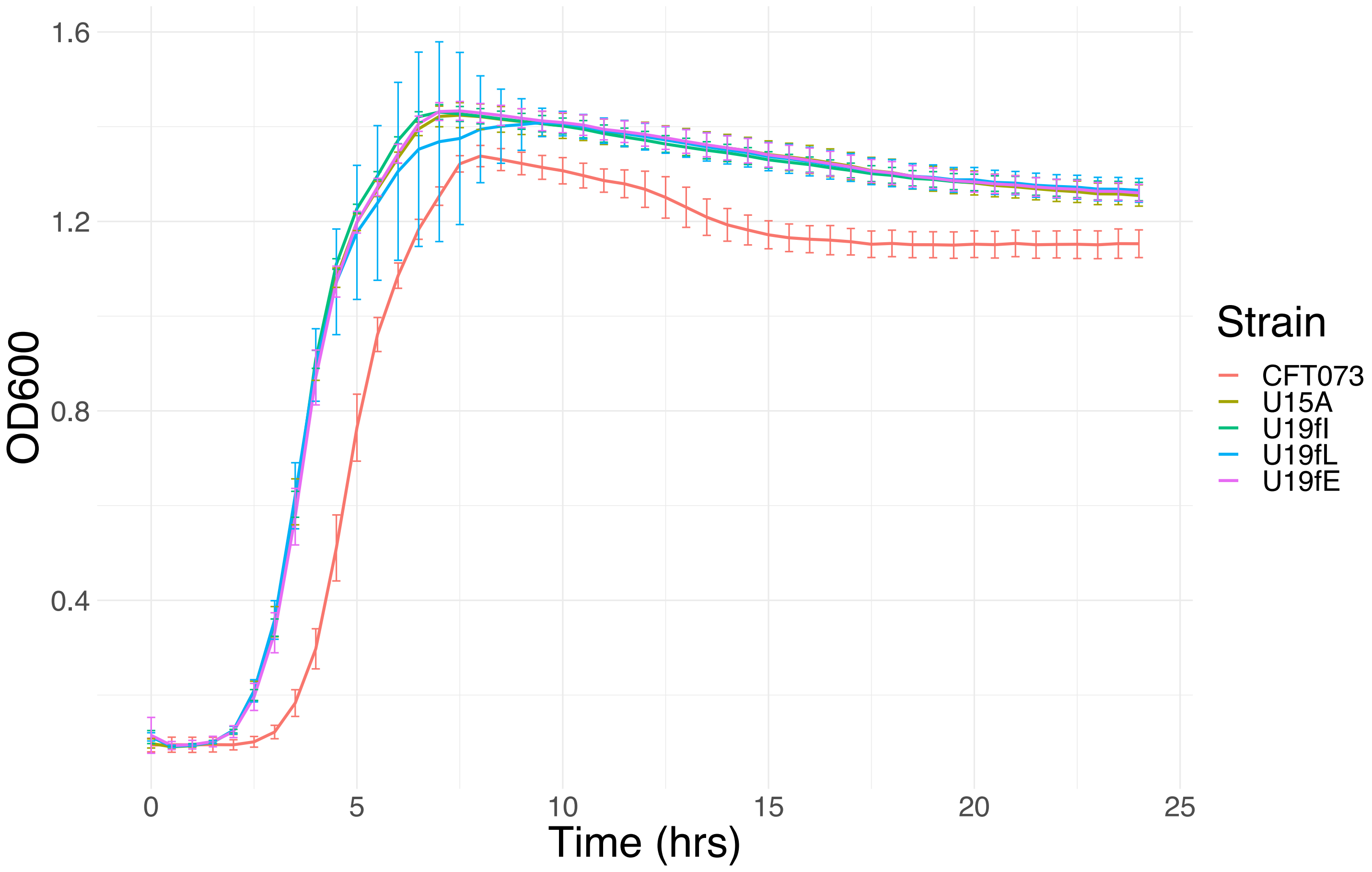


**Figure S5.** Growth curves of *E. coli* strains CFT073, U15A, U19tE, U19cL, and U19cI in LB broth.

**Table S8.** Genes encoded by strain U15A that are not encoded by strain U19fE.

| Genes | Genbank Accession or NCBI RefSeq | % Identity |
|---|---|---|
| Pertactin-like passenger domain-containing protein | AGY87489.1 | 100 |
| Conjugal transfer protein *traI* | HAM0114830.1 | 99.63 |
| Hypothetical | ELE89707.1 | 100 |
| *aac(3)-IId*, aminoglycoside 3-N-acetyltransferase | ESD46483.1 | 100 |
| Tyrosine-type recombinase/integrase | MCW1595794.1 | 100 |
| Conjugal transfer protein *traF* | WP_000976351.1 | 100 |
| Hypothetical | AGQ80681.1 | 100 |
| Hypothetical | ABE09421.1 | 100 |
| Histidine phosphatase family protein | WP_000977522.1 | 100 |
| Hypothetical | EQN57353.1 | 100 |
| Rep protein, partial | MDF8590270.1 | 100 |
| Replication protein | WP_227633736.1 | 100 |
| Hypothetical | ADT77573.1 | 100 |
| *tmrB,* tunicamycin resistance protein | HBN4657367.1 | 100 |
| Hypothetical | WP_212737122.1 | 100 |
| Conjugal transfer protein *traE* | WP_001238939.1 | 100 |
| Conjugal transfer lipoprotein *traH* | WP_001079808.1 | 100 |
| Hypothetical | ASK38484.1 | 100 |
| Hypothetical | UGG79343.1 | 100 |

**Table S9.** Genes encoded by strain U19cL that are not encoded by U19cI.

| Genes | Genbank or NCBI Accession | % Identity |
|---|---|---|
| Hypothetical | ART44382.1 | 100 |
| Hypothetical | AIX64318.1 | 100 |
| YehK protein/hypothetical | AAN81103.1 | 100 |
| *ilvB* operon leader peptide IvbL | WP_001312198.1 | 100 |
| Hypothetical | WP_001065661.1 | 100 |
| Hypothetical. Contains YgeY family selenium metabolism-linked hydrolase domain. | GCN70441.1 | 94.64 |
| Hypothetical | ABE07489.1 | 100 |
| Hypothetical | CEK06034.1 | 95.24 |

1. B. M. Forde *et al.*, Population dynamics of an *Escherichia coli* ST131 lineage during recurrent urinary tract infection. *Nat Commun* **10**, 3643 (2019).
